# Proteomics profiling of serum and liver in GSD Ia and Ib patients: insights into complication mechanisms and circulation biomarkers

**DOI:** 10.1101/2025.09.08.674441

**Authors:** Ruiqi Xiao, Candelas Gross-Valle, Albert Gerding, Adam M. Thorne, Maaike H. Oosterveer, Terry G.J. Derks, Vincent E. de Meijer, M. Rebecca Heiner-Fokkema, Barbara M. Bakker, Justina C. Wolters

## Abstract

**Background:** Glycogen Storage Disease (GSD) Types Ia and Ib are rare metabolic diseases caused by gene variants in *G6PC1* and *SLC37A4,* respectively. Although life-threatening fasting hypoglycemia can be controlled by a strict diet, patients often suffer from multiple metabolic abnormalities and severe long-term complications. However, the underlying mechanisms remain incompletely understood, and the lack of effective monitoring biomarkers makes it a challenge to treat patients. Therefore, the aims of this study are to investigate the pathological mechanisms of the disease and disease complications in GSD I and identify potential protein biomarkers.

**Methods:** In this study, we employed comprehensive untargeted proteomics on stored samples: 26 serum or plasma samples from 18 GSD Ia and 8 GSD Ib patients with 21 matched control sera, complemented by 4 liver samples from 2 GSD Ia patients who received liver transplantation (from the one patient with hepatocellular carcinoma we obtained tissue from both the carcinoma tissue and the adjacent non-carcinoma tissue), and 1 from a GSD Ib patient, compared to 10 donor liver samples.

**Results:** We identified a total of 415 proteins in our analyses. Pathway analysis of the differentially regulated proteins revealed distinct changes in serum/plasma of GSD Ia and Ib. The coagulation pathway was the most significantly changed biological process in the GSD Ia patients. Immune response-associated proteins, especially a large number of immunoglobulins, were increased in GSD Ib specifically. Proteins related to liver injury, cholesterol, and amyloidosis were altered in two subtypes, though more pronounced in GSD Ia. Potential biomarkers with significant alterations both in the circulation as well as in the liver tissue were identified specifically for monitoring GSD I subtypes and prognosing liver deterioration, namely GSD Ia (COL163 and PROC), GSD Ib (F11 and CD163), and hepatocellular carcinoma (HCC) in GSD Ia patients (ALDOB and CFHR5).

**Conclusions:** These findings provide new insights into the differences between the two GSD I subtypes and the pathogenesis of GSD I-related complications, as well as highlighting the potential of protein circulating biomarkers for monitoring complication progression in GSD I and assessing HCC risk in GSD Ia patients.

**Trial registration:** Not applicable.

**A concise 1 sentence take-home message (synopsis) of the article:** This comprehensive in-depth proteomics study on blood and liver GSD Ia and Ib patient samples provided novel insights into understanding of the disease subtypes as well as the the pathogenesis of GSD I-related complications, such as HCC risk in GSD Ia patients and identified potential biomarkers for the disease subtypes and complications.

## Background

Glycogen Storage Disease I (GSD I) is an inherited disease of carbohydrate metabolism.^1^ GSD Ia (OMIM: #232200) and Ib (OMIM: #232200) are subtypes of GSD I, which are caused by gene variants in *G6PC1* (OMIM: *613742)^2^ and *SLC37A4* (OMIM: *602671) ^3^, respectively, leading to dysfunction of the glucose 6-phosphatase enzyme (G6PC) or the glucose-6-phosphate transporter (G6PT).

GSD I patients suffer from severe fasting intolerance and hepatomegaly, and is biochemically characterized by hypoglycemia, hyperlipidemia, hyperlactatemia, and hyperuricemia.^4,5^ Although dietary management can help reduce part of these symptoms, patients are still at risk of acute and chronic complications, such as growth retardation, liver and kidney disease.^5,6^ Notably, in GSD Ia hepatocellular adenoma (HCA)^1^ and carcinoma (HCC)^7^ are more common, compared to GSD Ib. In addition, GSD Ib patients suffer from neutropenia and neutrophil dysfunction,^8,9^ increasing the risks of infections and inflammatory bowel diseases.^10^ Patients show significant clinical heterogeneity in the long-term complications, and the underlying mechanisms for these differences are only partially understood. Further understanding of these differences is crucial for treatment and monitoring, as well as for risk stratification of long-term complications.^3,11^ Several omics studies have been applied to gain a deeper understanding of GSD, which mainly have focused on GSD I (Table 1). ^12–26^ In GSD I, metabolic dysregulation was observed in glucose, fatty acids, amino acids, creatine, purine/pyrimidine, and lipids, despite dietary intervention.^19,21,24,26^ On the glycosylation level, hypo-galactosylation and accumulation of high-mannose and hybrid type *N-*glycans was observed in GSD Ib,^13,20^ and ApoC-III hypo-sialylation was reported in both GSD Ia and Ib.^16^

**Table 1.** Summary of the omics study in GSD.

| First Author | GSD Subtype | Sample species | Sample Details | Omics Method | Key Findings |
| --- | --- | --- | --- | --- | --- |
| Baodong Sun, et al., 2009 <sup>12</sup> | GSD-Ia | Mouse | Liver (3 glucose treated, 3 non-glucose treated GSD Ia, 3 controls) | Proteomics | Elevated GAPDH in G6Pase-deficient hepatocytes promoted apoptosis, shown by increased TUNEL positivity and caspase-3 activation in GSD-Ia livers. |
| Bu'Hussain Hayee, et al., 2011 <sup>13</sup> | GSD-1b | Human | neutrophils (5 G6PC3-deficient, 2 GSD Ib patients) | Glycomics | Neutrophil hypo-galactosylation of *N*- and *O*-glycans in G6PC3 deficiency and GSD Ib patients. |
| Julien Calderaro, et al., 2013 <sup>14</sup> | GSD-Ia, GSD-Ib | Human | Liver (25 HCA and 7 non-HCA frozen samples from 14 GSD Ia, 1 Ib) | DNA sequencing | GSD-related HCA characterized by a lack of HNF1A inactivation. High frequency of b-catenin mutations could be related to the increased frequency of malignant transformation in HCC. |
| Li-Ya Chiu, et al., 2014 <sup>15</sup> | GSD-Ia | Human | Liver (paired adenomas and normal liver tissues from 7 GSD Ia); Serum (6 GSD Ia, 20 controls) | miRNA sequencing | Distinct miRNA deregulation in HCA associated with GSD Ia, and miR-130b was a potential circulating biomarker for HCA. |
| Nina Ondruskova, et al., 2018 <sup>16</sup> | Types 0, Ia, non-Ia, III, IX | Human | serum (39 samples, 24 patients: 2 type 0, 7 Ia, 3 non-Ia, 7 III, 5 IX) | Glycomics | GSD types Ia and non-Ia exhibited mild ApoC-III hyposialylation, whereas types III and IX displayed severe glycosylation defects. |
| Davide Cangelosi, et al., 2019 <sup>17</sup> | GSD Ia | Mouse | Liver (20 GSD Ia vs. 18 WT) | Proteomics | Metabolic reprogramming, inflammation, and M2 macrophage polarization in GSD Ia mouse liver, potentially promoting tumorigenesis. |
| Roberta Resaz, et al., 2020 <sup>18</sup> | GSD Ia | Mouse | Plasma exosomes (45 GSD Ia mice vs. 18 WT) | MicroRNAs | Altered exosomal microRNA expression in GSD Ia mice, linked to glucose metabolism, inflammation, and hepatocellular carcinoma. |
| Luciana Hannibal, et al., 2020 <sup>19</sup> | GSD Ia, GSD Ib, GSD III | Human | Fibroblasts (1 GSD Ia, 1 Ib, 3 III vs. 1 healthy) | Metabolomics | Distinct metabolic signatures in GSD subtypes, including altered energy metabolism, oxidative stress response, and creatine metabolism. |
| Bobby G. Ng, et al., 2021 <sup>20</sup> | GSD Ib | Human | Serum (7 GSD Ib), induced pluripotent stem cell (iPSC)-derived hepatocytes from fibroblasts (1 GSD Ib), and CRISPR-edited Huh7 cells) | Glycomics | Dominant *SLC3744* mutation cause liver-specific congenital disorders of glycosylation (CDG) with coagulopathy, and accumulation of high mannose and hybrid type *N*-glycans in serum. |
| Tamara Mathis, et al., 2022 <sup>21</sup> | GSD Ia, GSD Ib | Human | Plasma (14 GSD I patients: 11 GSD Ia, 3 GSD Ib; 31 controls) | Metabolomics | Broad metabolic disturbances in plasma, including fuel metabolism, lipid metabolism, amino acid, methylation, urea cycle, and purine/pyrimidine metabolism. |
| Eran Tallis, et al., 2022 <sup>22</sup> | GSD Ib | Human | Plasma (1 GSD Ib patient, pre/post empagliflozin) | Metabolomics | Empagliflozin treatment reduces 1,5-anhydroglucitol, improves neutrophil function, and alleviates IBD symptoms in a GSD Ib patient. |
| Karina Colonetti, et al., 2023 <sup>23</sup> | GSD Ia, GSD Ib, GSD III/IX | Human | Plasma (16 GSD Ia, 6 Ib, 5 III/IX, 24 controls) | Multiplex luminex assay for 20 cytokines | Patients (GSD Ia/III/IX) presented reduced IL-4, MIP-1α/CCL3, et al. GSD Ib showed elevated G-CSF and IL-10. GSD III/IX had increased levels of IP-10/ CXCL10. IP10/CXCL10 and MCP-1 were higher in GSD III/IX. GSD I patients with anemia presented higher levels of IL-4 and MIP-1α. Triglycerides correlated with cytokine levels. |
| Alessandro Rossi, et al., 2024 <sup>24</sup> | GSD Ia | Human | Serum (12 GSD Ia vs. 13 healthy and 7 hypercholesterolemia controls) | Targeted lipidomics | Total cholesterol and triglyceride values correlated with ceramide (Cer). Circulating Cer may allow for refined dietary assessment compared with traditional biomarkers. |
| Yuxuan Lu, et al., 2025 <sup>25</sup> | GSD II (Pompe) | Human | Muscle (6 late-onset Pompe vs. 4 controls) | Proteomics + Metabolomics | 635 upregulated and 89 downregulated proteins; significant activation of mTOR, autophagy, and lysosome pathways; reduced L-arginine levels. |
| Francesca Pirozzi, et al., 2025 <sup>26</sup> | GSD Ia | Human | Serum (12 GSDIa vs. 12 controls) | Proteomics + Metabolomics | The altered levels of liver-specific proteins and choline-related metabolites in GSD Ia patients reveal liver damage and lipid dysregulation. ALDOB correlates with ALT and AST, supporting its potential as a biomarker for long-term monitoring of liver injury. |

On the proteome level, a targeted study screening for 20 cytokines was performed on GSD I patient material, showing altered inflammatory signatures for both GSD Ia (reduced IL-4) and GSD Ib (elevated IL-10).^23^ More in-depth screening of the protein adaptations was performed specifically on GSD Ia. ^26^ This study revealed liver-specific protein alterations associated with liver damage and lipid dysregulation, with the identification of up-regulated aldolase B as a potential biomarker for liver injury.^26^ Mouse studies on GSD Ia revealed extensive protein remodeling associated with metabolic reprogramming, including inflammatory pathway activation and M2 macrophage polarization, which potentially promote tumorigenesis.^17^ However, these proteomics studies did not include subjects with long-term complications. Current identification of biomarkers for long-term complications have so far only been achieved for HCA/HCC in GSD I by DNA and miRNA sequencing, identifying HNF1A (DNA)^14^ and miR-130b (miRNA)^15^ as potential biomarkers for HCA in GSD I.

Identification of reliable tumor marker is essential since the classical biomarkers for HCA and HCC using imaging and serum markers, including alpha-fetoprotein (AFP) and carcino-embryonic antigen (CEA), which were initially suggested for GSD I by 2002 European guidlines.^27^ However, in the more recent American College of Medical Genetics and Genomics (ACMG) guideline, AFP and CEA haven been shown to remain normal in GSD I patients with HCC.^6^ Different groups have reported the proteindes-gamma-carboxy prothrombin (DCP, or also called PIVKA-II) as a potential biomarker for HCC in GSD Ia more recently, but these observations are so far only described in four patients.^28–30^ Underlining the importance for identification of reliable protein biomarkers in large(r) patients cohorts.

Therefore, we performed an explorative comprehensive proteomics profiling in serum and plasma of GSD Ia and Ib patients, with a subset of patients had already developed HCA and HCC. The aims of this study are understanding the mechanism of the both GSD I subtypes and identifying prognostic biomarkers for GSD Ia and Ib patients, especially for HCC in GSD Ia patients.

## Methods

### Study participants

This study was conducted in accordance with the principles of the Declaration of Helsinki. For patients followed up at the University Medical Center Groningen (UMCG), the local Medical Ethics Committee determined that the Medical Research Involving Human Subjects Act did not apply to retrospective, non-interventional studies, and therefore, additional ethical approval was not required (METc 2019/119). The experimental protocols were reviewed and approved by the METc at UMCG, following the relevant guidelines and regulations outlined in protocol METc 2019/119.

This study enrolled blood (serum or plasma) and liver samples (Figure 1A). The details of those patient samples were summarized in Tables S1 and S2. Briefly, 18 GSD Ia serum or plasma samples were taken from 12 GSD Ia patients (age range 12-49 years; 6 females and 6 males; 9 non-HCA/HCC from 8 patients, 7 HCA from 3 patients, and 2 HCC from 1 patient). 8 GSD Ib serum samples were taken from 5 GSD Ib patients (age range 11-19 years, 3 females and 2 males; non-HCA/HCC), as well as 21 serum or plasma samples from age- and sex-matched individuals (age range 10-50 years; 10 females and 11 males) without metabolic diseases. All retrospective blood samples in this study were serum, with the exception of A10, C01, C03, C04, and C19 were plasma. In addition, 4 liver samples (L_A02_HCC of tumor tissue, L_A02 of non-tumor tissue, L_A08, and L_B05) were from 2 GSD Ia (A02 and A08) and 1 GSD Ib (B05) patients, with 10 age- and sex-matched control liver samples (6 adult controls and 4 pediatric controls) from liver transplantation donors (Figure 1A). Multiple samples from the same patient, collected at different times were labeled using ‘a’, ‘b’, and ‘c’ (e.g., A03a and A03b).

**Figure 1.**
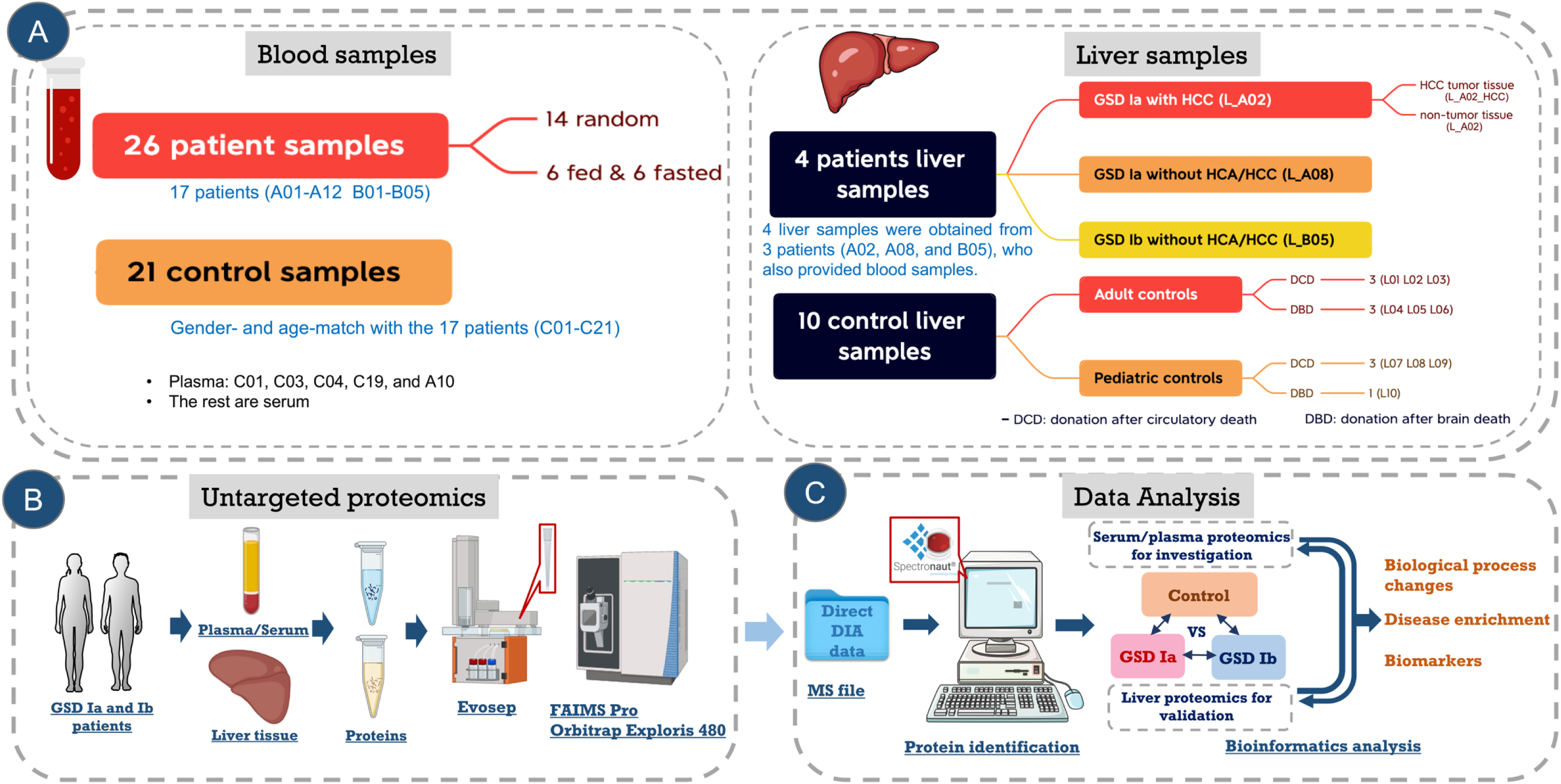
Summary of samples, untargeted proteomics workflow, and data analysis.

### Untargeted Proteomics workflow

An untargeted proteomics workflow was established to identify protein groups of serum/plasma and liver samples (Figure 1B), followed by data analysis (Figure 1C). Liver tissue proteomes were also included, providing a unique insight into the link between the liver and the circulation. The liver proteomics results were used to prioritize the identified potential plasma biomarkers by selecting proteins that show differential regulation in both sample types as the strongest indicators to assess the hepatic health status of these patients.

### Liver homogenization

10% lysis buffer (w/v, mg/mL) was added into liver tissues, a mechanical agitator (VWR, VOS PB S40) was used at 700 for 20 times to make liver homogenates. The lysis buffer was made by 48.45 mL 0.1% Nonidet P-40 buffer, 0.5 mL phosphatase inhibitors cocktail 2 (Sigma-Aldrich, P5726), 0.5 mL phosphatase inhibitors cocktail 3 (Sigma-Aldrich, P0044), 50 μL 1M DL-Dithiothreitol (DTT, Sigma-Aldrich, 43819), and one complete protease inhibitor cocktail tablet (Roche, 11836145001). 0.1% Nonidet P-40 buffer was made by 0.4 M sodium chloride (NaCl, Sigma-Aldrich, 746398), 1:1000 Nonidet P-40 (NP40, Merck KGaA, 492016), 10 mM Tris(hydroxymethyl)aminomethane (Duchefa, T1501.1000), and 1 mM ethylenediaminetetraacetic acid (EDTA, Sigma-Aldrich, E4884-100G). The primary liver homogenate was collected into 1.5 mL tubes and sonicated on ice (Sonics & Materials INC, VCX 130) two times for 10 seconds each, with pulse on/off 1 second. After sonicating, samples were centrifuged at 3000 rpm, at 4 °C for 10 mins, and the supernatant collected into new tubes as 10% liver homogenates.

### Delipidation

90 μL liver homogenates and 20 μL serum/plasma were delipidated using 4 mL MeOH and 4 mL cold diethyl ether (-20 °C), followed by centrifugation for 30 minutes at 1,500 xg at 4 °C. After discarding the supernatant, 4 mL cold diethyl ether was added to each sample and centrifuged for 30 minutes at 1,500 x g at 4 °C. Samples were dried under N_2_ after discarding the supernatant, and then resuspended with 100 μL 1× lithium dodecyl sulfate (LDS) sample loading buffer (Abcam PH=8.5, diluted with RIPA buffer). After sonicating, vortexing, pipetting up and down several times, and centrifugation at 1,500 xg for 1 min, the supernatants were taken and diluted 5 times using 1×LDS sample buffer.

### In-gel digestion

The in-gel digestion protocol was based on the ‘In-Gel Digestion and Sample Cleanup’ protocol, as described previously in Wolters et al.^31^ Briefly, 20 μL of delipidated samples were loaded on a precast polyacrylamide gel (Bolt™ Bis-Tris Plus Mini Protein Gels, 4-12%, 1.0 mm, Thermo Scientific) and ran for 5 minutes at 100V. The gels were stained for one hour with InstantBlue® Coomassie Protein Stain (Abcam), and after destaining by MiliQ water, the blue gel bands were cut into small squares of 2 mm×2 mm and collected separately. They were washed and shaken at 500 rpm on Thermomixer comfort with 300 μL 30% v/v acetonitrile (ACN) in 100 mM ammonium bicarbonate (ABC) until they were completely destained, followed by 300 μL 50% ACN in ABC, and lastly with 300 μL 100% ACN for 5 min, after which they were dried in the oven (Melag, brutschrank incubat) at 37 °C. Next, the proteins were reduced with 30 μL 10 mM dithiothreitol (30 minutes, 55 °C) and alkylated with 30 μL 55 mM iodoacetamide (30 min, RT, in the dark), and then they were shaken with 300 μL 100% ACN for 30 minutes, following by removed the solution and dried in the oven at 37 °C. Subsequently, the gel pieces were rehydrated with 30 μL trypsin (1:100 g/g, Sequencing Grade Modified Trypsin, V5111, Promega), and incubated overnight at 37 °C. The residual liquid was collected in a new tube the next day, and the peptides in the gel plugs were extracted in 30 μL 75% v/v ACN plus 5% v/v formic acid (FA) (20 min, 25 °C, mixing 500 rpm). After combining these two liquid fractions, peptides were dried in a vacuum concentrator (Eppendorf). The residues were resuspended in 0.1% FA and diluted to 25 ng/μL. Further analysis was standardized based on the protein concentration before delipidation.

### Sample loading and liquid chromatographic separation

Twenty μL peptide solutions (25 ng/μL) were loaded onto the Evotips (Evosep Aps, Odense C, Denmark) following the sample loading protocol,^32^ which were ready for the liquid chromatograph-mass spectrometer (LC-MS) analysis. All samples were analyzed by the Evosep One HPLC (Evosep Biosystems, Odense C, Denmark) using a C18 column with length/ID/C18 bead size of 15 cm/150 μm/1.9 μm (EV-1106, Evosep Biosystems). The pre-programmed gradient (30 samples per day, 0.5 μL/min flow rate, 48 minutes cycle time, 44 minutes gradient length) was applied. The column temperature was maintained at 40 °C by a column heater controller (Phoenix S&T, Chadds Ford, USA) and interfaced online with a FAIMS Pro, and then to the Orbitrap Exploris 480.

### Mass spectrometric analysis

Mass spectrometric analyses were performed on Orbitrap Exploris 480 (Thermo Scientific) using a HRMS1-DIA based-method^33^ coupled to a FAIMS Pro (Thermo Scientific). The FAIMS Pro was set to a standard resolution with 3.8 L/min total carrier gas flow and voltages switching between, -45 and -60 CV. MS^1^ and MS^2^ data were acquired at a resolution of 120k and 15k, across a 400-1,200 m/z and 400-1,000 m/z scan range, and with normalized AGC targets of 300% and 1,000%, respectively. MS^1^ and MS^2^ were acquired with a maximum inject time of 400 ms. HRMS1-DIA settings included 3 full scans and 35 fragment scan events with 17 m/z isolation windows, 32% HCD collision energies, 40% RF lens, and 12 loop spectra.

### Data analysis and statistics

Raw MS data were processed in Spectronaut v.19.4.241120.62635 using a library-free approach (directDIA) for quantification at MS^1^ level, using background signal as the imputation strategy. Data was aligned with protein sequences based on the Human UniProt reviewed sequences (20422 entries). The mass spectrometry proteomics data have been deposited to the ProteomeXchange Consortium via the PRIDE ^34^ partner repository with the dataset identifier PXD066805. The gene names related to the proteins were used for visualization and their integrated peak areas were processed using RStudio. The proteins with missing values of more than 50% across all samples were removed from further analysis based on the published guidance.^35^ The rest of the missing values were imputed by the mean of the respective experimental group. If no group values were available, missing values were replaced by 0.1, representing the instrument detection limit. ‘EdgeR’ package in R Studio v.4.2.2^36^ was used to normalize data and calculate log2^Fold Change^ (Log2^FC^) and false discovery rates (FDR) between groups. In addition, principal component analysis (PCA), heatmap, volcano plots, and barplots were made using R packages ‘sjmisc’, ‘ggrepel’, ‘pheatmap’, and ‘ggplot2’. Significantly changed proteins (SCPs) were filtered by FDR (≤ 0.05).

Gene Ontology (GO) analysis and DisGeNET disease enrichment analysis were performed using DAVID Bioinformatics Resources (v2024q2). In the disease enrichment results, we excluded the proteins related to dysfibrinogenemia because 5 plasma samples and 42 serum samples were employed, because a previous study described that the fibrinogen proteins are highly abundant in plasma, which could introduce bias with mixed sample types.^37^ The GO and disease enrichment results were visualized as bar plots using the R package ‘ggplot2’. For biomarkers identification, if there were at least two samples in each comparison group, R package ‘pROC’ was utilized to calculate the area under the curve (AUC) and make the receiver operating characteristic (ROC) curves of proteins with AUC>0.9.^38^ In the liver tissue, given that only one sample was available for GSD Ia with HCC and for the GSD Ib liver, proteins were considered potential biomarkers if their intensities were higher than 1.2 times the maximum or lower than 0.8 times the minimum observed in the reference group.

## Results

### GSD Ia and Ib show different serum/plasma protein patterns

An overview of untargeted proteomics shows the alterations in GSD I samples (Figure 2). 21 Controls, 18 GSD Ia (including 7 HCA, 2 HCC, and 9 without HCA/HCC), and 8 GSD Ib samples were grouped. A total of 415 proteins were identified and mapped on a heatmap to identify any potential clustering across all samples (Figure 2A). Principal component analysis (PCA) further revealed clear separation between groups, suggesting substantial global protein differences among GSD Ia, GSD Ib, and control samples (Figure 2B). GSD Ia patients with HCA and HCC clustered together (Figures 2B and S1B), suggesting additional specific changes in the proteome in relation to these long-term complications. The potential confounding factors, including age, gender, sample type, and feeding state (fed, fasted, random), did not show clustering in the PCA analyses (Figures S1C-S1F). The volcano plots show 158 (38%), 116 (28%), and 151 (36%) proteins were significantly changed in the GSD Ia vs. Control, GSD Ib vs. Control, and GSD Ia vs. Ib comparisons (Figure 2C-2E and Table S3-S5). Overall, these results provided a comprehensive overview and indicate clear differences in proteins not only between GSD I and controls, but also between the GSD Ia and Ib subtypes, and between non-HCA/HCC and HCA/HCA GSD Ia patients.

**Figure 2.**
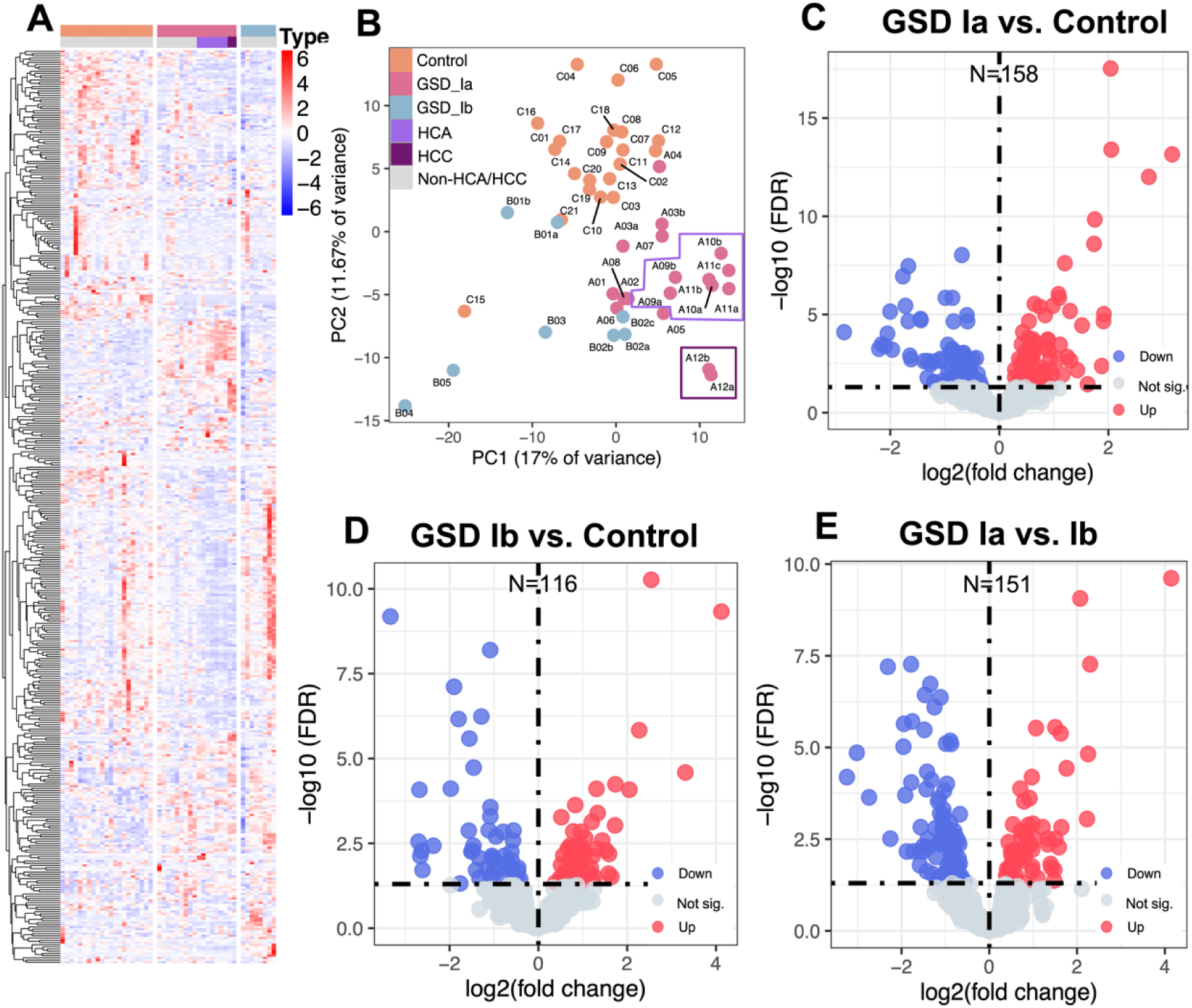
Overview of serum/plasma proteomics analysis across GSD Ia, GSD Ib and control groups. (A) Heatmap showing the z-score of all proteins detected in all serum/plasma samples from all groups. (B) Principal Component Analysis (PCA) plot illustrates the proteomic profile separation among groups. (C-E) Volcano plots of differential protein between groups. Significantly up-regulated and down-regulated proteins (FDR < 0.05) are highlighted in red and blue, respectively. ‘HCA’ represents GSD Ia patients with Hepatocellular adenoma (HCA), and ‘HCC’ refers to those who initially had HCA and later progressed to hepatocellular carcinoma (HCC). ‘N’ refers to the number of significantly changed proteins (SCPs), including ‘up’ and ‘down’ proteins. ‘Not sig.’ refers to proteins that show no significant changes. Multiple samples from the same patient, collected at different times were labeled using ‘a’, ‘b’, and ‘c’ (e.g., A03a and A03b).

### Biological processes related to immune response, blood coagulation, and complement activation are the primarily affected processes in serum/plasma of GSD Ia and Ib

To understand the context and biological relevance of these differences between the three groups, the SCPs in different comparisons (GSD Ia vs. Control, GSD Ib vs. Control, and GSD Ia vs. Ib) were clustered according to the Gene Ontology biological process (Table S6-S8) to identify the main biological pathways that were altered in the circulation of the GSD I patients. Out of the top 10 p-value-ranked differentially regulated biological processes for each of the three comparisons between GSD Ia, GSD Ib, and control showed overlap in the immunoglobulin mediated immune response, blood coagulation, complement activation (classical pathways), and fibrinolysis (Figure 3A). The immune response pathways showed the most significant alterations, with the top 3 altered pathways in both GSD Ib vs. Control (blue bars) and GSD Ia vs. Ib (yellow bars) being immune-related and showing considerable overlap. GSD Ia vs. Control (pink bars) showed differences in immunoglobulin mediated immune response, blood coagulation, and complement activation as the top 3 changed pathways. This suggests that these pathways are consistently affected in patients but exhibit subtype-specific variations in their alterations.

**Figure 3.**
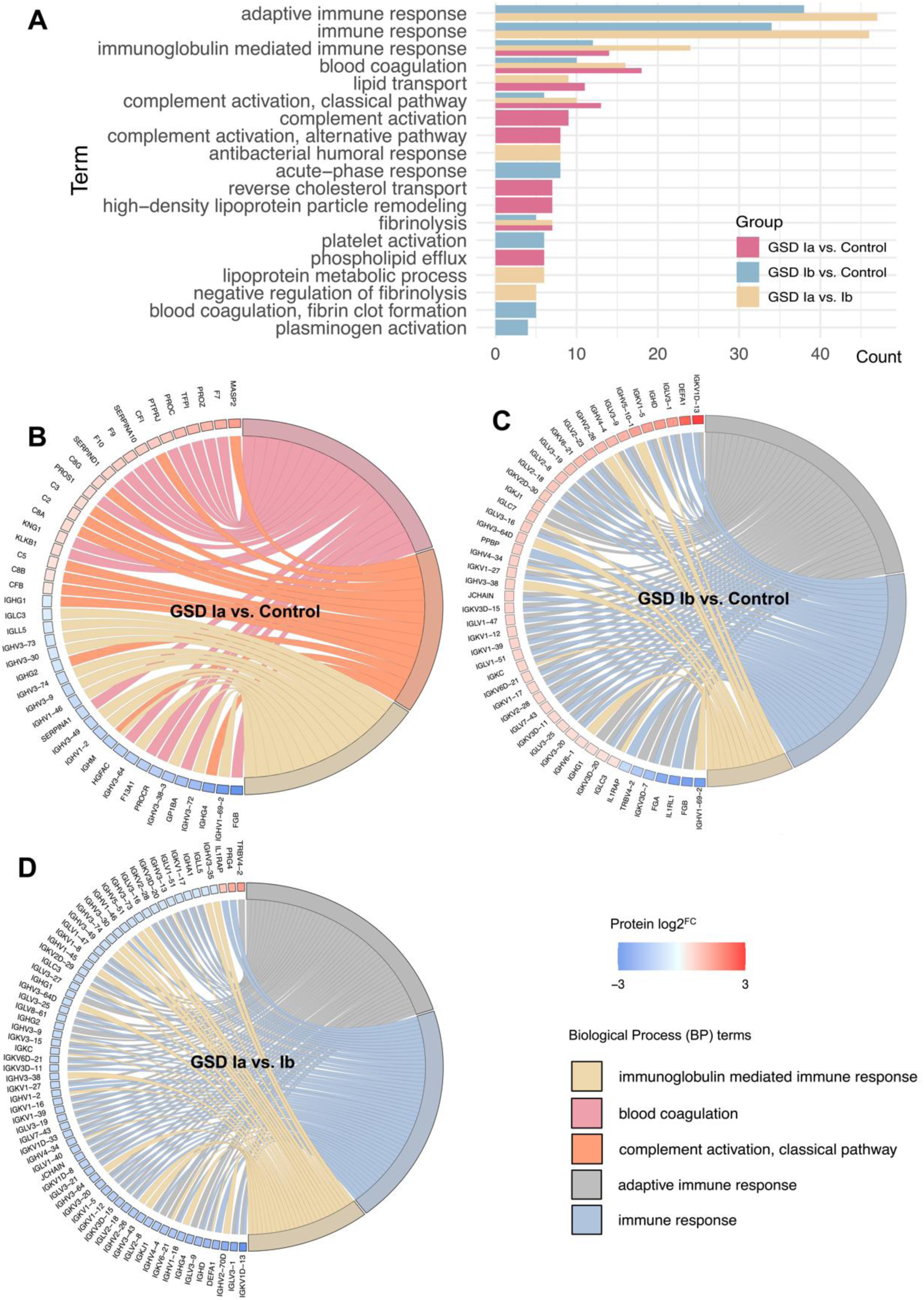
Gene Ontology (GO) biology process (BP) analysis of SCPs in serum/plasma among Control, GSD Ia, and GSD Ib groups. (A) Bar plot showing the BP pathways enriched across SCPs in all three comparisons. The bar length reflects the number of SCPs associated with each pathway. (B-D) Chord plots illustrate the top three enriched BP pathways for (B) GSD Ia vs. Control, (C) GSD Ib vs. Control, and (D) GSD Ia vs. Ib. Each plot highlights the pathways and the specific SCPs contributing to enrichment, with connections representing the relationships between proteins and pathways.

To visualize the specific proteins contributing to these pathway differences, we created chord diagrams of the top 3 enriched pathways for each comparison (Figure 3B-3D). In GSD Ia vs. Control, proteins were mainly up-regulated (marked with red boxes) in blood coagulation (12/18, 67%) and complement activation pathways (9/13, 69%), while all proteins were down-regulated (marked with blue boxes) in the immune response (14/14, 100%) (Figure 3B). Notably, 17 immunoglobulins (IGs) were decreased in GSD Ia (Figure 3B), while 95% of the IGs were increased in GSD Ib (38/40, except IGHV1-69-2 and IGKV3D-7) (Figure 3C). This difference was also clear from the comparison between GSD Ia and Ib (Figure 3D), where IGs (62/67) were also the proteins providing the largest differentiation between the two subtypes.

Taken together, the blood coagulation and complement activation pathways were the top 2 changes in GSD Ia, and the immune response pathway changed both in GSD Ia and Ib. Most of the immune response proteins were decreased in GSD Ia vs. Control, while increased in GSD Ib vs. Control, and we observed that these opposite effects provide the largest differentiation between GSD Ia vs. Ib. The top 3 altered biological processes in GSD Ia vs. Ib were all related to the immune response pathway. These differences in the types and trends of altered proteins between GSD Ia and Ib result in distinct coagulation and immune disruptions.

### Significantly changed proteins in GSD I are primarily related to disease clusters: liver injury, thrombosis, complement and antibody deficiency, hypercholesterolemia and hyperlipoproteinemia, and amyloidosis

The context of proteins can alternatively be viewed from the predicted risks for disease complications, using a disease enrichment analysis (DisGeNET) on the serum/plasma SCPs from these three comparisons ((GSD Ia vs. Control, GSD Ib vs. Control, and GSD Ia vs. Ib) to identify protein- and pathway-disease associations (Tables S9-S11). The bar plots (Figure 4A) show the top 10 ranked disease clusters from the DisGeNET results, which were grouped in five overarching topics based on their function (Figure 4B). Namely, liver injury, thrombosis, complement and antibody deficiency, hypercholesterolemia and hyperlipoproteinemia, and amyloidosis.

**Figure 4.**
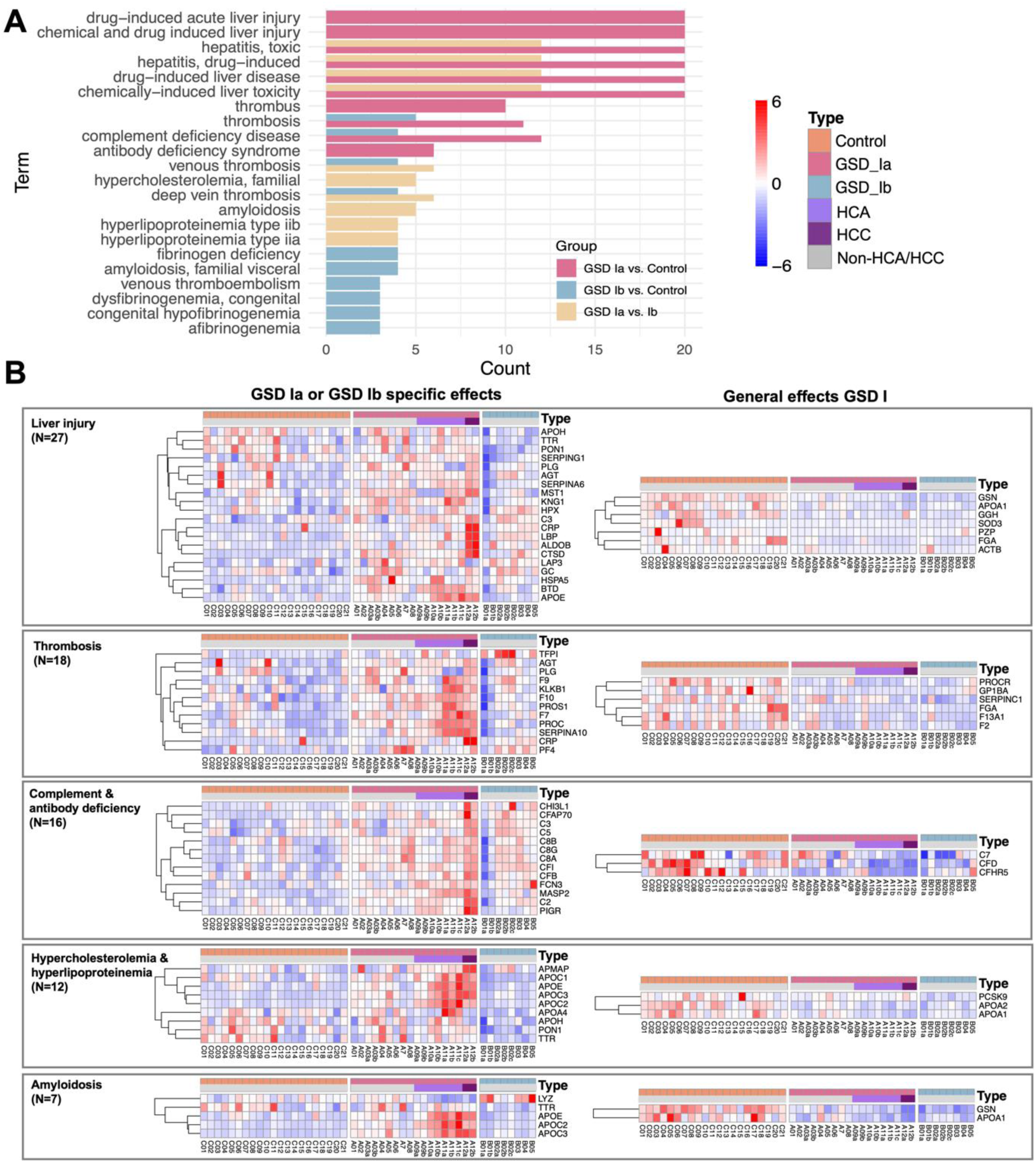
Disease cluster analysis of SCPs in serum/plasma. (A) Barplot showing the enrichment of SCPs in various diseases. (B) Heatmaps represent the distribution of SCPs across different disease categories, as classified from panel A, highlighting the interesting proteins involved in each category. ‘N’ represents the number of SCPs.

In the identified overarching disease clusters, proteins were divided into two groups, either as a specific effect for GSD Ia or GSD Ib or as a general effect observed in both GSD I subtypes (Figure 4B). All the proteins with general effects in GSD Ia and Ib showed similar down-regulation when compared with controls. Notably, in the ‘GSD Ia or GSD Ib specific effects’ group, the majority of the proteins showed only significant up-regulation in GSD Ia in all these clusters, when compared to controls. In the ‘General effects GSD I’ group, F2, a key protein in blood clot formation, was decreased in both GSD Ia and Ib within the thrombosis-related cluster (Figure 4), consistent with the altered blood coagulation pathway identified in both subtypes (Figure 3A). Interestingly, the apolipoproteins showed distinct mixed changes: APOA1 and APOA2 were decreased in both subtypes; APOA4 was decreased in GSD Ib; while APOC1, APOC2, APOC3, and APOE were increased in GSD Ia.

These analyses provided a focus on additional proteins involved in long-term complication processes, next to the immunoglobulin and coagulation pathways that were primarily identified in the GO clustering analyses.

### Potential prognostic biomarkers in blood circulation were identified specific for GSD Ia (COL6A3 and PROC) and GSD Ib (F11 and CD163)

To identify potential biomarkers, we applied ROC curve analyses for all the significantly altered proteins in the different comparisons in serum/plasma and liver (Figure S2). In the serum/plasma, 23 potential protein biomarkers were found in GSD Ia vs. Control group (Figure 5A), 36 in GSD Ib vs. Control (Figure 5B), and 51 in GSD Ia vs. Ib (Figure 5C).

**Figure 5.**
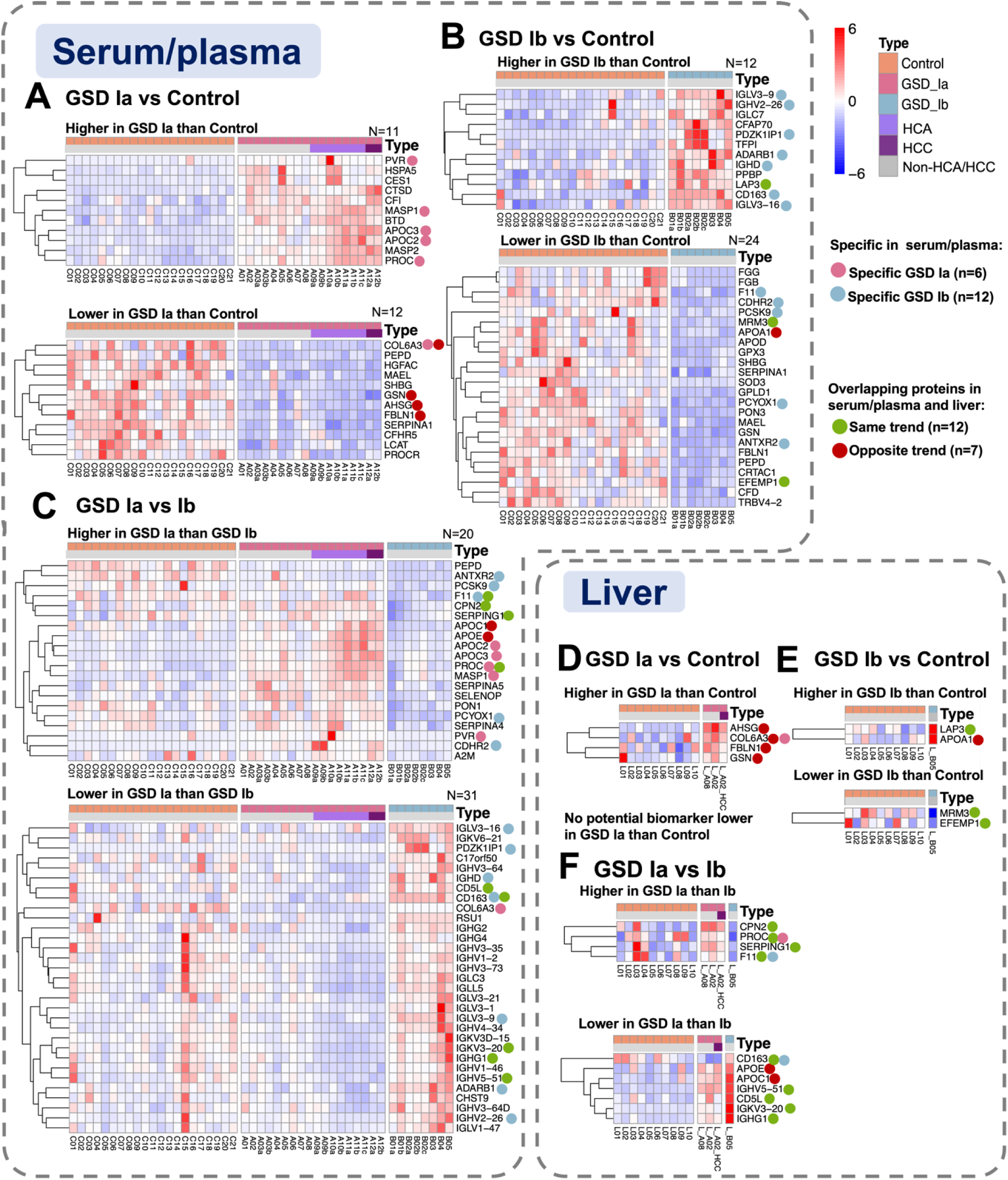
Summary of potential biomarkers for GSD Ia and Ib, when compared with Control and each other in serum/plasma and liver. Heatmap showing the z-score of (A-C) potential protein biomarkers detected in serum/plasma samples, and (D-F) overlapping protein biomarkers in the liver. ‘N’ refers to the number of proteins.

In these selections, we further highlighted the subset of potential biomarkers that could uniquely identify the specific GSD subtype, by annotating the targets that were specific for either GSD Ia (Figure 5A and 5C, pink dots n=6) or GSD Ib (Figure 5B and 5C, blue dots n=12) in comparison to both control and the other GSD subtype.

Alternatively, we used our unique liver tissue proteomics data to further focus our selection of potential serum protein biomarkers, by identifying the proteins that were significantly changed both in serum/plasma as well as in the liver proteome. This yielded a total of 4 potential biomarkers for GSD Ia vs. Control (Figure 5D), 4 proteins for GSD Ib vs. Control (Figure 5E) and 11 proteins for GSD Ia vs. Ib (Figure 5F). The information and direction of the changes in serum/plasma and liver are summarized in Table 2. Four of these proteins were also identified before as subtype specific serum/plasma biomarkers, namely COL6A3 (down-regulated in serum/plasma, up-regulated in liver) and PROC (up-regulated in both serum/plasma and liver) specific for GSD Ia, and F11 (down-regulated in both serum/plasma and liver) and CD163 (up-regulated in both serum/plasma and liver) as specific biomarkers for GSD Ib vs. Control (highlighted with pink and blue dot in Figure 5D-5F).

**Table 2.** Summary of the potential biomarkers of different comparisons. ’↑’ refers to increase, ‘↓’ refers to decrease. Proteins highlighted in pink and blue are specific biomarkers for GSD Ia and GSD Ib, respectively.

| Biomarkers for | Protein Name | Trend in serum/plasma | Trend in liver | Functional Category | potential related complications in GSD I | Description |
| --- | --- | --- | --- | --- | --- | --- |
| **GSD Ia vs. Control** | AHSG | ↓ | ↑ | Acute phase protein / Immune regulation | Immune disorder | Alpha-2-HS-glycoprotein (Fetuin-A), regulates calcium metabolism and immune response; a negative acute-phase protein. |
|  | **COL6A3** | ↓ | ↑ | Extracellular matrix / Structural protein | Liver injury | Collagen type VI alpha 3 chain, part of extracellular matrix structure, maintains tissue integrity. |
|  | FBLN1 | ↓ | ↑ | Extracellular matrix / Structural protein | Liver injury | Fibulin-1, involved in cell adhesion and stabilization of the extracellular matrix. |
|  | GSN | ↓ | ↑ | Cytoskeletal regulatory protein | Liver injury | Gelsolin, modulates actin remodeling, associated with cell migration and apoptosis. |
| **GSD Ib vs. Control** | LAP3 | ↑ | ↑ | Enzyme / Proteolysis | Liver injury | Leucine aminopeptidase, involved in intracellular protein degradation and antigen processing. |
|  | APOA1 | ↓ | ↑ | Lipid metabolism | Hyperlipoproteinemia | Major component of HDL, responsible for reverse cholesterol transport. |
|  | MRM3 | ↓ | ↓ | Mitochondrial enzyme / Transcription regulation | Liver injury | Mitochondrial rRNA methyltransferase, involved in mitochondrial protein synthesis regulation. |
|  | EFEMP1 | ↓ | ↓ | Extracellular matrix / Structural protein | Liver injury | Also known as Fibulin-3, involved in ECM structure and regulation of cell proliferation. |
| GSD Ia<br>vs.<br>GSD Ib | CPN2 | ↑ | ↑ | Coagulation regulation / Inflammation | Coagulation disorder, Inflammation, Immune disorder | Carboxypeptidase N subunit, degrades peptide hormones and regulates inflammatory mediators. |
|  | PROC | ↑ | ↑ | Coagulation regulation | Coagulation disorder, | Protein C, an anticoagulant that inhibits coagulation cascade. |
|  | SERPING1 | ↑ | ↑ | Complement inhibition / Inflammation | Coagulation disorder, Inflammation, Immune disorder | C1 inhibitor, controls complement activation and prevents excessive inflammation. |
|  | F11 | ↑ | ↑ | Coagulation factor | Coagulation disorder, | Coagulation factor XI, involved in the intrinsic coagulation pathway. |
|  | CD163 | ↓ | ↓ | Macrophage marker / Immune regulation | Immune disorder | Clears hemoglobin-haptoglobin complexes; anti-inflammatory macrophage marker. |
|  | APOE | ↑ | ↓ | Lipid metabolism | Hyperlipoproteinemia | Regulates cholesterol transport, linked to atherosclerosis and neurodegenerative diseases. |
|  | APOC1 | ↑ | ↓ | Lipid metabolism | Hyperlipoproteinemia | Modulates lipoprotein metabolism, affects VLDL and LDL balance. |
|  | IGHV5-51 | ↓ | ↓ | Immunoglobulin variable region / Immune response | Immune disorder | Immunoglobulin heavy chain variable region gene, contributes to antibody diversity. |
|  | CD5L | ↓ | ↓ | Immune regulation / Lipid metabolism | Immune disorder, Hyperlipoproteinemia | Also known as AIM, regulates lipid metabolism and macrophage function. |
| **GSD Ia vs. GSD Ib** | IGKV3-20 | ↓ | ↓ | Immunoglobulin variable region / Immune response | Immune disorder | Immunoglobulin light chain variable region, involved in antigen recognition. |
|  | IGHG1 | ↓ | ↓ | Immunoglobulin constant region / Immune effector | Immune disorder | IgG1 subclass, mediates antibody-dependent cytotoxicity and complement activation. |
| **Specific GSD Ia with HCC** | ALDOB | ↑ | ↑ | Carbohydrate metabolism enzyme | Liver injury | Aldolase B, key enzyme in fructose metabolism, primarily expressed in the liver. |
|  | CFHR5 | ↓ | ↓ | Complement regulation | Immune disorder | Complement factor H-related protein, modulates the alternative complement pathway to protect host tissue. |

### ALDOB and CFHR5 were identified as potential prognostic biomarkers in GSD Ia with HCC

The subset of GSD Ia patients with HCA and HCC provided us with the unique opportunity to identify potential prognostic biomarkers for the HCA/HCC long-term complications (Figure S3). In total, 33 proteins were identified as potential prognostic markers for tumors in GSD Ia, as they already show significant alterations when the HCA and HCC patients were combined as one group (HCA+HCC), compared to non-HCA/HCC GSD Ia and Controls (Figure 6A). Further focusing specifically on HCC (compared to all other samples) yielded a total of 84 proteins as potential biomarkers for monitoring of HCC in GSD Ia in serum/plasma (Figure 6B).

**Figure 6.**
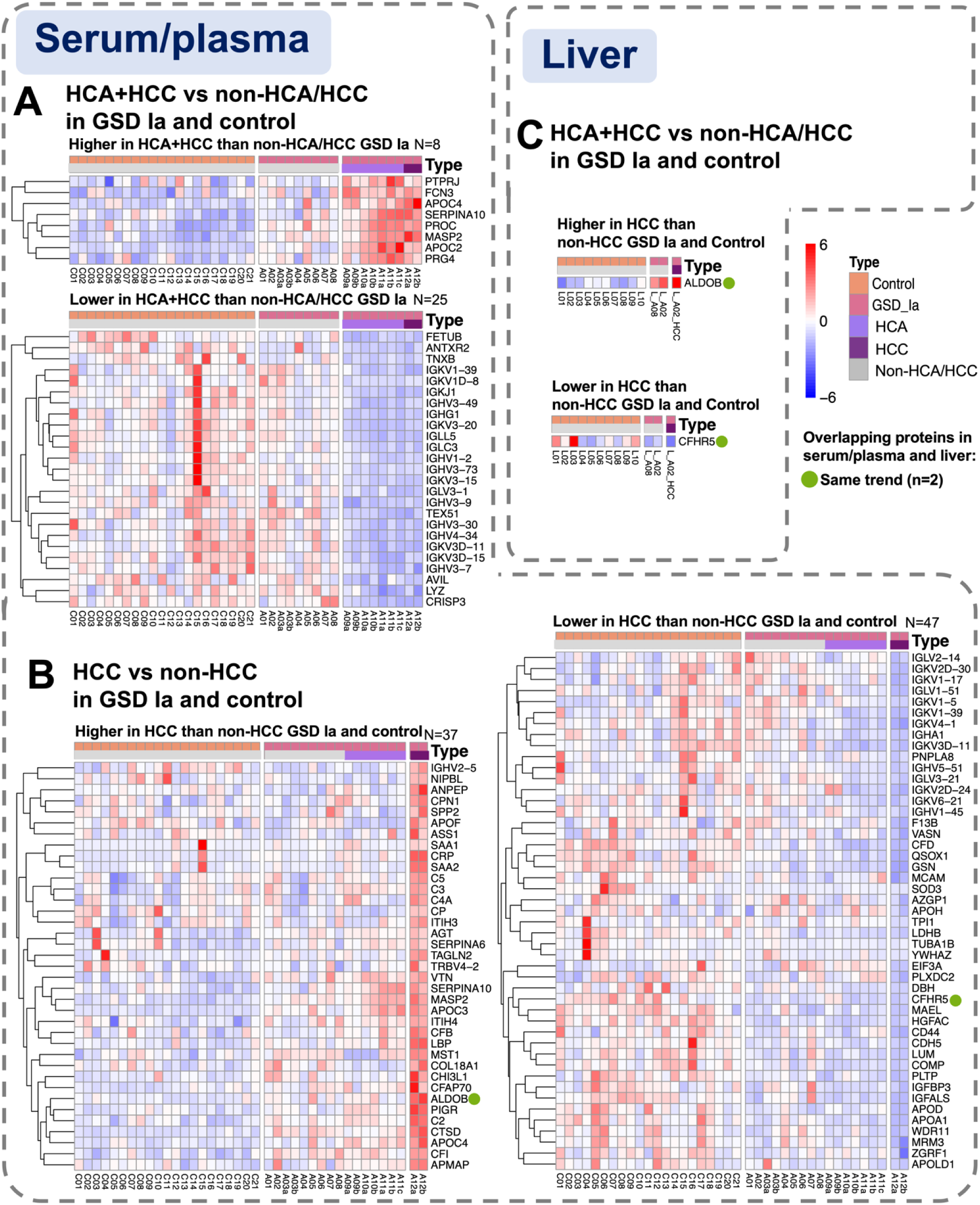
Summary of potential biomarkers for HCA+HCC and HCC in serum/plasma and liver. Heatmap showing the z-score of (A and B) potential protein biomarkers detected in serum/plasma samples, and (C) overlapping protein biomarkers in the liver. ‘N’ refers to the number of proteins.

We also used our liver proteomics data to identify the subset of targets also showing significant changes in the liver for further selection of interesting HCC biomarkers found in serum/plasma. This analysis highlighted two proteins, namely ALDOB (up-regulated) and CFHR5 (down-regulated), both showing the same directionality in serum/plasma and in the liver proteomics (Figure 6C), identifying them as potential key biomarkers for HCC in GSD Ia.

In summary, all the identified biomarker candidates for GSD Ia, GSD Ib and the role of HCA/HCC in GSD Ia were listed in Table 2, including protein trends in serum/plasma and liver, functional categories and descriptions. They were related to immune regulation, cellular structure, cytoskeletal regulation, lipid metabolism, transcription regulation, coagulation regulation, inflammation, and carbohydrate metabolism.

## Discussion

Previous GSD omics studies have characterized metabolic changes and identified some mRNA, metabolic, and protein biomarkers (Table 1)^12–26^. Our research systematically analyzed the untargeted proteomics of samples from both GSD Ia and Ib patients in a relatively large cohort. Two elements made this study unique in design: the inclusion of patients who had already developed HCA or HCC, and the correlation of circulating proteomics markers with the target tissue (i.e., the liver). Our results provide new insights into GSD Ia- and Ib-specific proteomic alterations and their possible links to biological processes and disease complications, providing leads for monitoring of the GSD I subtypes and prognostic biomarkers.

Our pathway analyses identified blood coagulation as the most significantly altered process in GSD Ia (Figure 3A), which is connected to the thrombosis disease cluster (Figure 4B). This could potentially link with clinical findings that a coagulation dysfunction has been observed in GSD I patients, who have easy bruising, epistaxis, and excessive bleeding during surgery.^39–42^ Additionally, a previous study in our group by La Rose et al. indicated that hepatocyte-specific glucose-6-phosphatase deficiency disturbs platelet aggregation and decreases blood monocytes upon fasting-induced hypoglycemia in GSD Ia mouse models.^43^ In their study, the plasma levels of coagulation factors V (FV) and VIII (FVIII) were increased in GSD Ia mice in the fed and fasted conditions, while factor VII (FVII) was not affected. In line with their study, we validated that the G6PC deficiency disturbs coagulation in GSD I patients. In our study, 18 proteins involved in thrombosis were altered in GSD Ia and Ib patients, with 6 proteins decreased in both subtypes, especially the decrease of coagulation factor II (F2, also known as prothrombin). Prothrombin is the precursor of thrombin (F2a), the enzyme that provides the cleavage reaction to create the fibrin required for blood clot formation.^44–46^ Thus, as a key factor in coagulation, the decrease of F2 could provide a monitoring biomarker for the risk of coagulation complications in GSD I patients. Notably, F2 represents the fully γ-carboxylated prothrombin, whereas PIVKA-II, elevated in the serum of four GSD Ia patients with HCC,^28–30^ corresponds to the undercarboxylated form. In our results, F2 was increased in GSD Ia patients without HCC rather than in those with HCC, suggesting that different forms of prothrombin might reflect distinct disease states. Whether a shift from the fully γ-carboxylated to the undercarboxylated form contributes to disease progression, particularly during HCC, remains unclear. Further investigations are needed in light of existing knowledge on prothrombin metabolism. The additional number of up-regulated proteins related to thrombosis specific for GSD Ia could provide novel leads for the clinical observation that coagulation complications are more common in GSD Ia than Ib.^47–49^

The top 3 altered biological processes in both GSD Ib, compared to controls, as well as GSD Ia were all related to immune responses, with the proteins being up-regulated in GSD Ib in comparison to the other two groups. In terms of clinical symptoms, GSD Ib patients specifically suffer from immune-related complications such as recurrent infection, chronic inflammation, and inflammatory bowel disease, due to neutrophil dysfunction.^10,50,51^ In our study, a large number of IGs were increased in serum/plasma of GSD Ib patients, suggesting their infection and inflammation are more severe or they are continuously in a proinflammatory state, providing leads for the distinct immunological phenotype of GSD Ib patients.^1,10^

Our disease enrichment analysis identified interesting clusters related to liver injury, hypercholesterolemia and hyperlipoproteinemia, and amyloidosis as potential biomarkers for GSD I. These clusters contain a small subset of proteins being down-regulated in both GSD Ia and Ib, while a number of proteins are specificly up-regulated only in GSD Ia, which could provide leads to why GSD Ia tends to show more pronounced liver long-term complications than GSD Ib.^10, 52, 53^ Notably, the observed protein changes, especially apolipoproteins related to hypercholesterolemia and hyperlipoproteinemia, is in line with previously describe alteration in lipid metabolites in GSD Ia and Ib patient plasms samples.^21^ The consistent down-regulation of APOA1 and APOA2 in both subtypes reflected a shared impairment in the formation of high-density lipoprotein (HDL) and chylomicrons,^54,55^ which may contribute to the dyslipidemia frequently reported in GSD I patients. The decrease of APOA4 in GSD Ib further indicated dyslipidemia, as APOA4 is essential for plasma chylomicron assembly.^56^ It is known that GSD Ia patients have increased lipids (triglycerides and cholesterol) levels, consistent with observed elevation of APOC1, APOC2, APOC3, and APOE in GSD Ia, given these proteins and lipids are integral components of lipoproteins. ^57–60^ APOC2 activates lipoprotein lipase, ^61^ which contribute to triglyceride clearance, whereas APOC3 and APOC1 inhibit it.^62,63^ APOE is a key ligand for remnant lipoprotein clearance, may further reflect disordered lipid clearance and increased lipoprotein turnover. ^64^ Therefore, the complex effects of altered apolipoproteins may underlie the impaired lipid metabolism observed in GSD Ia and Ib. In particular, the increase of APOCs and APOE in GSD Ia could potentially reflect the more severe hyperlipidemia, aligning with results of mouse experiments ^65^ and clinical observations in patients.^27,66^ In addition, previous studies also indicated that the VLDL metabolism was impaired in hepatic-GSD Ia mouse model,^65^ in agreement with the observed high VLDL level in GSD Ia patients,^67^ which we observed in the VLDL related APOE and APOC proteins.

ROC curve analyses provided us with a range of potential biomarkers, but the GSD I liver tissue data provided us with an additional selection criterium to define a shortlist of potential biomarkers with a clear link to the liver changes of these patients. This resulted in the shortlist of potential liver-linked biomarkers showing differential regulation not only in the circulation, but also in the liver (Table 2) for the various comparisons between GSD Ia, GSD Ib and controls.

From these blood circulation biomarkers, two were subtype specific for GSD Ia (COL6A3 and PROC), two were subtype specific for GSD Ib (F11 and CD163). COL6A3 encodes the α3 chain of type VI collagen, known to be increased in NAFLD ^68^ and HCC ^69^, which reflects ongoing fibrogenesis and tissue remodeling. The up-regulated hepatic COL6A3 observed in our study may thus indicate progressive chronic hepatic disease in GSD Ia because extracellular matrix remodeling of hepatocytes. Interestingly, it was down-regulated in blood circulation, which might result from reduced release or increased sequestration in ECM structures, indicating localized remodeling rather than systemic turnover. Thus, the decreased level of COL6A3 in GSD Ia patients may reflect the risk of liver complications.

To monitor coagulation-related complications, we identified two potential biomarkers with subtype-specific relevance, PROC for GSD Ia and F11 for GSD. The protease PROC (protein C), plays an important role in regulating anticoagulation^70^ and was identified as a potential prognostic biomarker for liver cirrhosis ^71^ and inflammation ^72^ . Our finding of down-regulated F11 (coagulation factor XI) can be linked with earlier observations that GSD I patients often have bleeding tendencies,^11,39–42^ suggesting that reduced F11 could contribute to the impaired coagulation function. ^73^ Because previous studies have not specifically addressed F11 levels in GSD I so far, a potential functional link between the observed down-regulation of F2 (coagulation factor II or prothrombin) and F11 is an interesting lead for follow-up studies, as both are essential components of the coagulation cascade.

The last identified potential biomarker for GSD Ib is scavenger receptor cysteine-rich type 1 protein M130 (CD163), an acute phase-regulated receptor involved in clearance and endocytosis of hemoglobin/haptoglobin complexes by macrophages and may thereby protect tissues from free hemoglobin-mediated oxidative damage.^74,75^ The observed elevated CD163 in both the circulation and liver of GSD Ib could provide a potential biomarker for systemic and local macrophage activation and chronic inflammation status, ^76^ highlighting its potential as a biomarker for inflammation and immune activation. These potential biomarkers provided novel insights into the subtype specificity of GSD I as well as the prognosis of complications for GSD Ia and Ib patients.

Lastly, we also identified prognostic biomarkers for HCA and HCC in GSD Ia patients. Previous studies indicated 70-80% of GSD Ia patients would suffer from HCA over 25 years old, and 10% of them progress to HCC.^6,53,77^ Thus, it is essential to identify circulating biomarkers for early detection and intervention of HCA and HCC in these patients. Due to the lack of liver material of GSD Ia patients with HCA we were not able to narrow this list of 33 potential early prognostic biomarkers, but initial focus for further validation could focus on the eight up-regulated proteins, since it is easier to detect up-regulated proteins and this selection does include the above identified GSD Ia specific protein biomarker PROC.

For HCC we had liver tissue material available to prioritize the selection, and we identified ALDOB (up-regulated) and CFHR5 (down-regulated) as the two most interesting candidates in both serum/plasma and liver proteomics. ALDOB, as a key enzyme in fructose metabolism, was identified up-regulated both in a GSD Ia mouse study ^78^ and was also identified as a potential biomarker for GSD Ia liver injury in the recently published serum proteomics screening on GSD Ia patients^26^. However, we show with the distinction between GSD Ia patients with and without HCC, which are most strongly detected in the case of long-term liver complications, further supporting the role as a potential prognostic biomarker specifically for GSD Ia specific liver injury. Notably, previous studies reported that ALDOB was decreased in HCC of (non-GSD) patients,^79,80^ It is also not known how these observations relate to the strictly limited fructose intake^81^ of the GSD I patients. In addition, the decrease of CFHR5, a plasma glycoprotein produced in the liver,^82^ can be a risk factor for HCC in GSD Ia patients based on our study. However, previous studies have indicated that CFHR5 levels increase in various malignancies, particularly during metastasis.^83–86^ We therefore see the same opposite trends for both biomarkers in GSD Ia compared to the general HCC patients. Suggesting that metabolic changes in GSD Ia patients themselves may also affect the metabolism of liver cancer cells, which might be a unique metabolic adaptation in GSD Ia patients, which is further supported by the previous observation in the lack of response for the general HCC marker AFP.^6^ Targeted measurements are needed to confirm the clinical significance and elucidate the precise roles of ALDOB and CFHR5 in GSD-related HCC and their potential as prognostic biomarkers for liver-specific long-term complications.

However, several limitations should be acknowledged. First, despite the relatively large sample size, it is sufficient to capture the full heterogeneity of GSD I patients, especially the liver tissue samples. Second, patients were on different treatment regiments, while the controls not. Thus, we cannot be ruled out that the identified proteins mark treatment rather than disease responses. Third, serum/plasma and liver samples were retrospectively collected, and probably not under the same conditions. Fourth, both serum and plasma samples were used, which may also have contributed to heterogeneity, although we did not see distinct clustering between both types (Figure S1D) and excluded the proteins known to be different between plasma and serum.^37^ Altogether, the differential treatment regimens and sampling conditions may have contributed to heterogeneity in outcomes. Therefore, although potential biomarkers were identified in our study, their clinical applicability, stability, and specificity require further validation through large-scale and prospective studies.

## Conclusions

In summary, this study was the first to employ untargeted proteomics on serum/plasma of both GSD Ia and GSD Ib patients and on liver tissue of the same patients (subset) compared to matched controls, providing valuable leads into the complications mechanisms and patient monitoring: (1) We first identified the pathways changes and disease enrichment for the monitoring of the GSD I subtypes. (2) We further focused the list of potential biomarkers from the pathway and disease cluster analyses for potential biomarkers showing differential regulation both in the circulation as well as in the liver (Table 2); From this list we identified specific subsets for the monitoring and prognostic biomarkers specific for GSD Ia (COL6A3 and PROC), GSD Ib (F11 and CD163) and HCC in GSD Ia (ALDOB and CFHR5).

## List of abbreviations

ABC: ammonium bicarbonate
CAN: acetonitrile
AUC: area under the curve
CD163: scavenger receptor cysteine-rich type 1 protein M130
DTT: DL-Dithiothreitol
EDTA: ethylenediaminetetraacetic acid
F11: coagulation factor XI
F2: prothrombin FA: formic acid
FDR: false discovery rates
G6PC: glucose 6-phosphatase
G6PT: glucose 6-phosphate transporter
GO: Gene Ontology
GSD Ia: Glycogen Storage Disease type Ia
GSD Ib: Glycogen Storage Disease type Ib
HCA: hepatocellular adenoma
HCC: hepatocellular carcinoma
HDL: high-density lipoprotein
IBD: inflammatory bowel disease
LDS: lithium dodecyl sulfate
NP-40: Nonidet P-40
PCA: principal component analysis
PROC: protein C
ROC: receiver operating characteristic
UMCG: University Medical Center Groningen

## Declarations

### Ethics approval and consent to participate

The data were obtained by experiment and data analysis. The tenets of the Declaration of Helsinki were followed. The Medical Ethical Committee of the University Medical Center Groningen (UMCG) stated that for retrospective, non-interventional studies the Medical Research Involving Human Subjects Act was not applicable and that further study approval by the Medical Ethical Committee was not required (METc 2019/119).

### Consent for publication

Not applicable

### Availability of data materials

Raw mass spectrometry data used for proteomic analysis are available through ProteomeXchange with the dataset identifier PXD066805. The other data that supports the findings of this study are available in the supplementary materials.

### Authors’ contributions

Ruiqi Xiao: Methodology, Sample preparation, Proteomics measurements, Data analysis, Manuscript. Candelas G. Valle: Biology information of GSD Ia patients from clinical, Writing-reviewing and Editing. Albert Gerding: Help with methodology and sample preparation, Writing-reviewing and Editing. M. Rebecca Heiner-Fokkema: Provide some of the serum samples of the control group, Writing-reviewing and Editing. Adam M. Thorne and Vincent E. de Meijer: Provide liver samples of the control group, Writing-reviewing and Editing. Terry G.J. Derks: Biological information of GSD Ia patients from a clinical perspective, Writing-reviewing and Editing. Maaike H. Oosterveer, Barbara M. Bakker, and Justina C. Wolters: Conceptualization, Supervision, Writing-reviewing, and Editing.

## Acknowledgments

The authors gratefully acknowledge the support from the Chinese Scholarship Council (CSC) (No. 202106220094) and De Cock-Hadders Foundation (No. 816719) for Ruiqi Xiao, United for Metabolic Diseases and Metakids (UMD-CG-2023-037) for Justina Clarinda Wolters. Figures were partly created using Biorender. We thank Fabian Peeks, Aycha Bleeker and Trijnie Bos for excellent technical assistance.

