## Supplementary material for "Proteomics profiling of serum and liver in GSD Ia and Ib patients: insights into complication mechanisms and circulation biomarkers": Figure S1

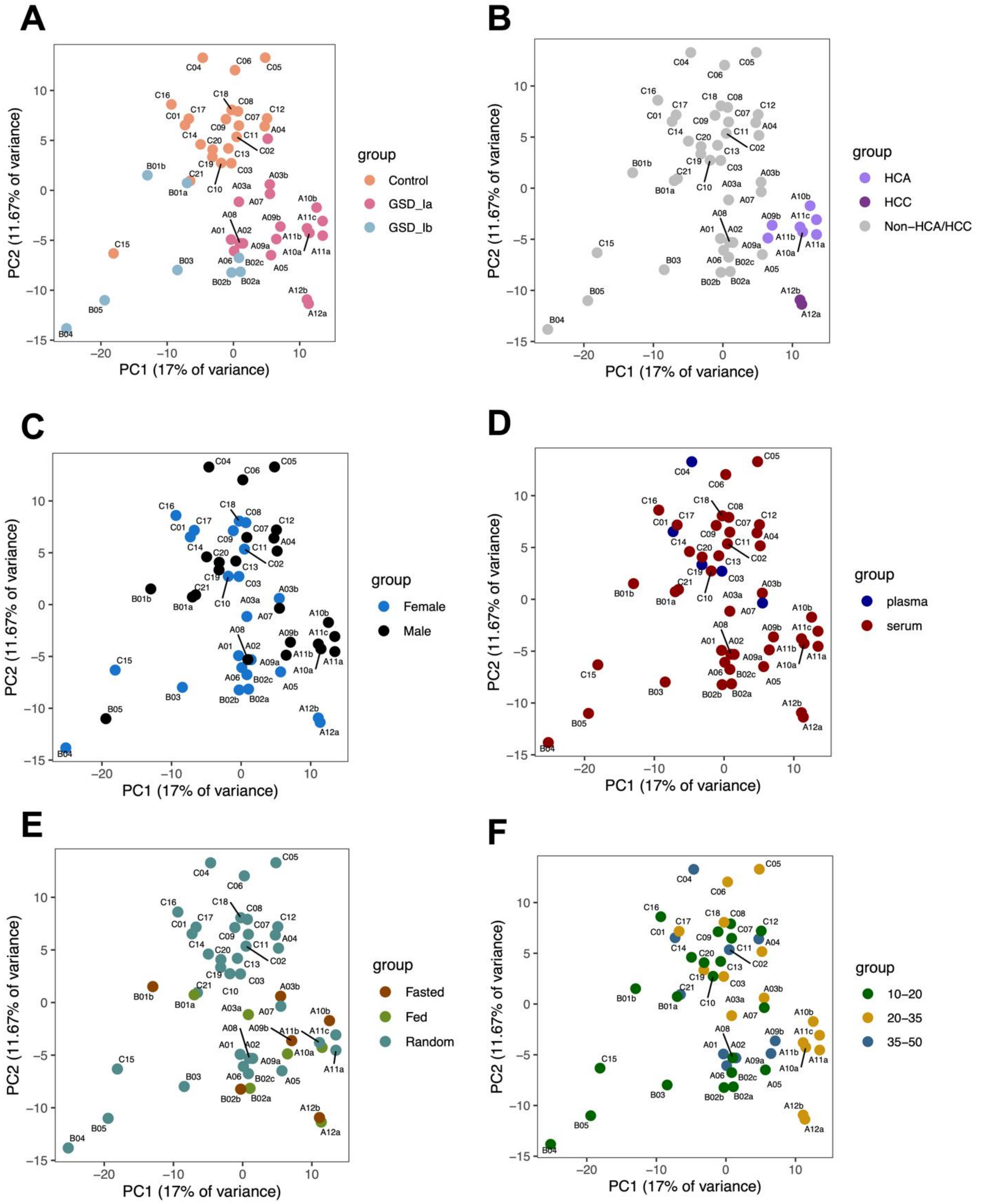


Figure S1. The Principal Component Analysis (PCA) plot illustrates the proteomic profile separation among groups. Multiple samples from the same patient, collected at different times were labeled using 'a', 'b', and 'c' (e.g., A03a and A03b).

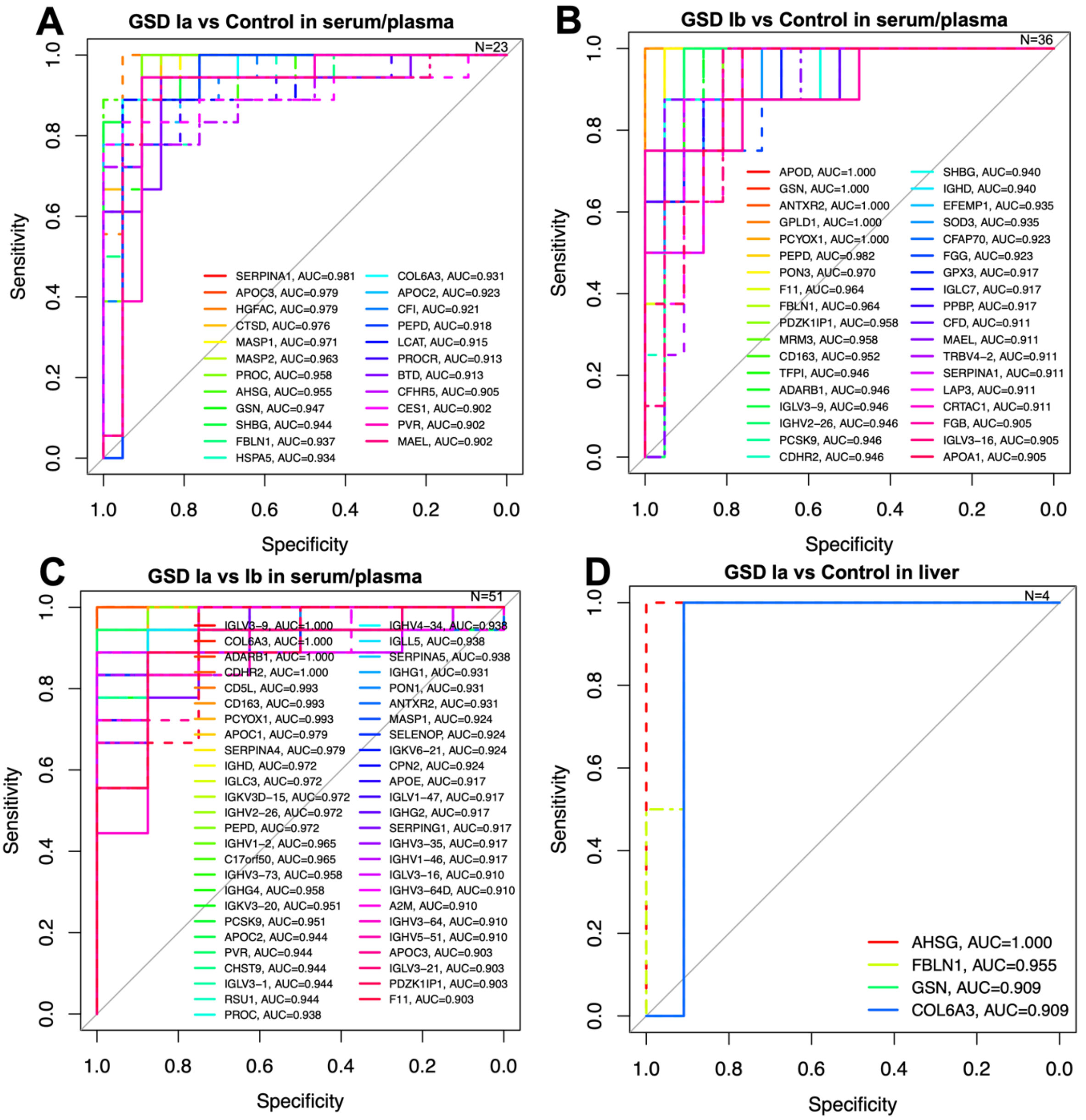


Figure S2. Receiver operating characteristic (ROC) curves of proteins with RUC>0.9 in: (A) GSD Ia versus Control, (B) GSD Ib versus Control, (C) GSD Ia versus GSD Ib in serum/plasma, and (D) GSD Ia versus Control in liver tissue. 'N' indicates the number of proteins with AUC>0.9.

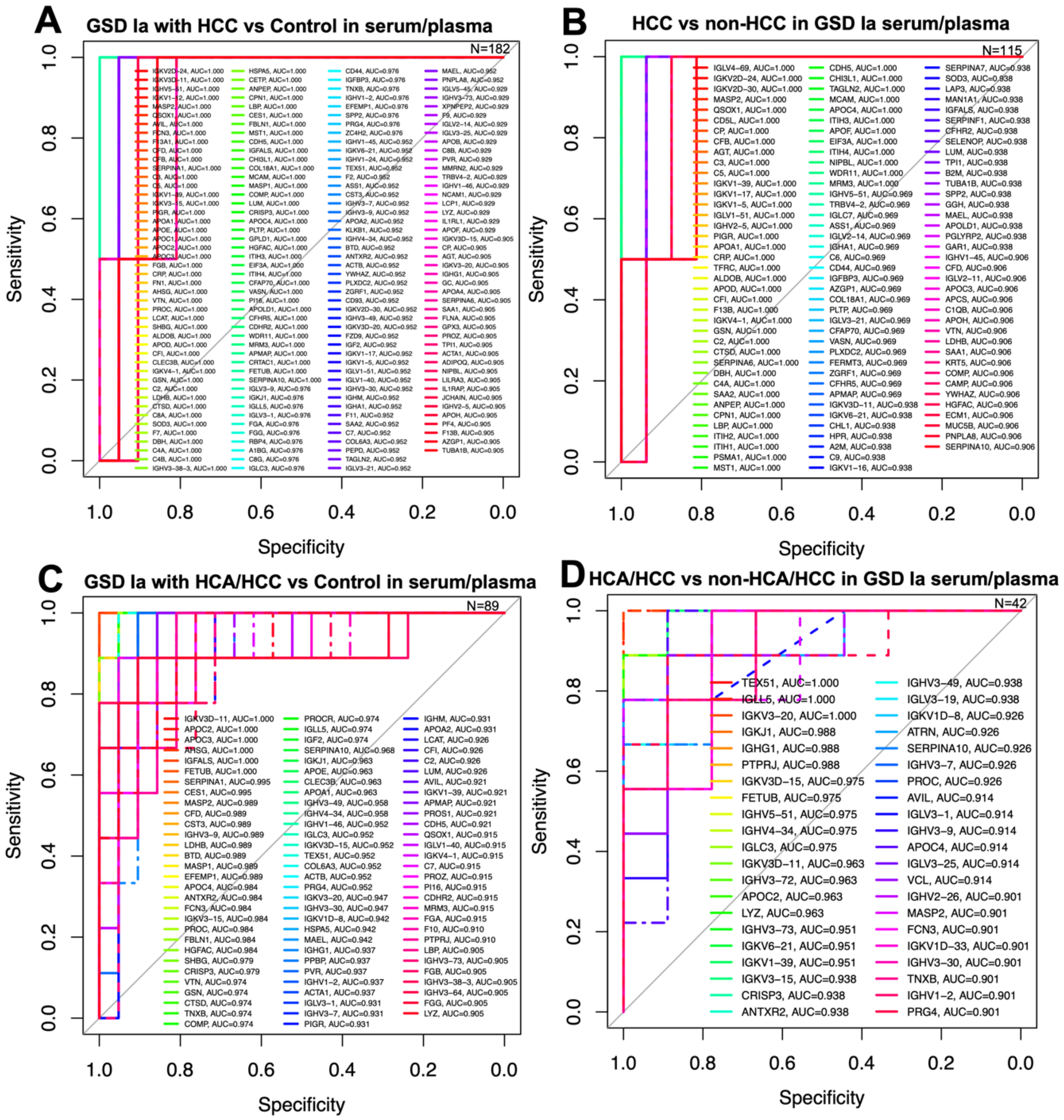


Figure S3. Receiver operating characteristic (ROC) curves of proteins with RUC>0.9 in: (A) GSD Ia with HCC versus Control, (B) GSD Ia with HCC versus those without HCC, (C) GSD Ia with HCA/HCC versus Control, and (D) GSD Ia with HCA/HCC versus those without HCA/HCC, based on serum/plasma analysis serum/plasma. 'N' indicates the number of proteins with AUC>0.9.
